# Single cell profiling of del(5q) MDS unveils its transcriptional landscape and the impact of lenalidomide

**DOI:** 10.1101/2023.10.19.562875

**Authors:** Guillermo Serrano, Nerea Berastegui, Aintzane Díaz-Mazkiaran, Paula García-Olloqui, Sofia Huerga-Dominguez, Ana Alfonso-Pierola, Marina Ainciburu, Amaia Vilas-Zornoza, Patxi San Martin, Paula Aguirre-Ruiz, Asier Ullate-Agote, Beñat Ariceta, Jose Lamo de Espinosa, Pamela Acha, Oriol Calvete, Tamara Jimenez, Antonieta Molero, Julia Montoro, Maria Díez-Campelo, David Valcarcel, Francisco Solé, Idoia Ochoa, Felipe Prósper, Teresa Ezponda, Mikel Hernaez

## Abstract

While del(5q) MDS patients comprise a well-defined hematological subgroup, the molecular basis underlying its origin, and the reason behind the relapse after lenalidomide remains unknown. Using scRNA-seq on CD34^+^ progenitor cells from patients with del(5q) MDS we were able to identify cells harboring the deletion, enabling us to deeply characterize the transcriptional impact of this genetic insult on disease pathogenesis and treatment response. We found, across all patients, an enrichment of del(5q) cells in GMP and megakaryocyte-erythroid progenitors not described to date. Interestingly, both del(5q) and non-del(5q) cells presented similar transcriptional lesions when compared to progenitors from healthy individuals, indicating that all cells, and not only those harboring the deletion, are altered in these patients and may contribute to aberrant hematopoietic differentiation. However, GRN analysis revealed a group of regulons with aberrant activity in del(5q) cells that could be responsible for triggering altered hematopoiesis, pointing to a more prominent role of these cells in the phenotype of these patients. An analysis of del(5q) MDS patients achieving hematological response upon lenalidomide treatment showed that the drug reverted several transcriptional alterations in both del(5q) and non-del(5q) cells, but other lesions remained, which may be responsible for potential future relapses. Moreover, lack of hematological response was associated with the inability of lenalidomide to reverse transcriptional alterations. Collectively, this study provides a deep characterization of del(5q) and non-del(5q) cells at single-cell resolution, revealing previously unknown transcriptional alterations that could contribute to disease pathogenesis, or lack of responsiveness to lenalidomide.

**KEY POINTS:**

– Del(5q) and non-del(5q) CD34+ cells share similar transcriptional alterations, with del(5q) cells presenting additional lesions.
– Hematological response to lenalidomide is associated with the reversal of some transcriptional lesions in del(5q) and non-del(5q) cells

## INTRODUCTION

Deletion of the long arm of chromosome 5 (del(5q)) is the most frequently observed cytogenetic alteration in *de novo* myelodysplastic syndromes (MDS). It affects around 10-15% of MDS patients and represents a distinct hematological and pathological subgroup due to its unique clinical features, such as macrocytosis, anemia, normal or high platelet count, and hypolobulated megakaryocytes^1^. With the advent of high-throughput sequencing technologies, two commonly deleted regions (CDR) were identified and mapped to del(5q): the 1.5 megabase region at 5q32-q33, which contains 41 coding genes^2,3^, and the 5q31 region containing 45 coding genes. Thus, the characterization of the molecular pathogenesis of del(5q) MDS has mainly focused on the identification of genes within these CDRs that are involved in the pathophysiology of the disease, such as *RPS14*, *miR-145*, *miR-146* and *EGR1*^4–7^. However, the presence of gene regulatory networks (GRNs) altered in these patients, as well as the transcriptional profile of hematopoietic progenitors from patients with del(5q), have not been analyzed in detail, in part due to the difficulty of identifying cells with and without the deletion within the same patient.

Lenalidomide represents the first therapeutic approach for del(5q) MDS, and induces in these patients a prolonged red blood cell transfusion independence and cytogenetic response^8^. Interestingly, even at the time of complete clinical and cytogenetic remission, persistent del(5q) progenitor cells have been identified, providing a reasonable explanation for the loss of responsiveness that patients experience in a 2-to 3-year interval^9^. However, the real impact of the deletion during lenalidomide treatment remains unknown, hinting the need of characterizing the molecular mechanisms driven by the deletion that could be impairing treatment response.

Previous studies have demonstrated that transcriptional alterations play a key role in MDS pathogenesis^10–14^. For example, our group recently identified the transcription factor *DDIT3* as a key erythropoietic regulator that is overexpressed in MDS, and showed its potential as a therapeutic target^15^. Furthermore, gene expression profiles have shown to be affected by different types of alterations, including cytogenetic abnormalities and mutations, among other factors^16,17^. In this sense, several studies have performed gene expression analyses in del(5q) MDS, and have identified several pathways deregulated in the disease, including those related to Wnt/β catenin or integrin signaling, as well as genes that could be potentially contributing to disease pathogenesis, such as *SPARC*^12,18^. Other studies have depicted the effect of lenalidomide on the gene expression profile of del(5q) patients, evidencing its potential to restore erythroid differentiation, modulate the bone marrow microenvironment, and revert the aberrant expression of putative pathogenic microRNAs^19–22^. However, the hematopoietic system of these patients is composed of a mixture of cells with and without the deletion (termed del(5q) and non-del(5q) cells, respectively), which masks the expression profile of the del(5q) cells, and limits the ability to define the transcriptional impact of the deletion.

In this work, we performed single-cell RNA-sequencing (scRNA-seq) in primary CD34^+^ hematopoietic progenitor cells from del(5q) MDS patients at diagnosis and patients after lenalidomide treatment, and applied copy number alteration (CNA) analyses to link the del(5q) genotype to the transcriptional profile of each individual cell. This approach yielded a well-characterized del(5q) MDS atlas. Leveraging the generated atlas, we detected previously unknown transcriptional alterations both in del(5q) cells and also in non-del(5q) cells at diagnosis and after lenalidomide treatment. We demonstrate that non-del(5q) cells present aberrant behavior compared to the healthy hematopoietic system similar to that observed in del(5q) cells. We also show that although lenalidomide restores some of the detected alterations of del(5q) and non-del(5q) cells of responder patients, other lesions identified at diagnosis remain after treatment. Furthermore, our results evidence that lenalidomide is not able to reverse part of the transcriptional lesions carried by del(5q) cells of a non-responder patient, which seems to be associated with the lack of hematological response.

## METHODS

### Sample collection

The samples and data from the patients included in the study were provided by the Biobank of the University of Navarra and were processed according to standard operating procedures. Patients and healthy donors provided informed consent, and the study was approved by the Clinical Research Ethics Committee of the Clinica Universidad de Navarra. Bone marrow aspirates were obtained from healthy elderly controls [(n=3), median age, 72 years, range, 61-84 years] and from patients with MDS [(n=7), median age, 84 years, range, 80-91 years] from the Clinica Universidad de Navarra and collaborating hospitals. The patients’ clinical characteristics are shown in Table S1.

### Fluorescence-activated cell sorting

For purification of CD34^+^ cells, BM samples were lysed in 1X of BULK lysis buffer (150 mM NH4Cl, 10 mM CHKO3 and 0.1 mM EDTA in deionized water) for 15 mins at a sample to bulk ratio of 1:10, and centrifuged for 5 mins at 1500 rpm to eliminate red blood cells. Next, cells were stained using CD34-APC (clone 581; Beckman Coulter) and CD45-PerCPCy5.5 (clone HI30; Biolegend) for 15 min at RT. CD34^+^ CD45^+^ cells were then sorted in a BD FACSAria II (BD Biosciences) and directly used for scRNA seq analysis.

### scRNA-seq library preparation

The transcriptome of the bone marrow CD34^+^ cells were examined using NEXTGEM Single Cell 3’ Reagent Kits v3 and v3.1 (10X Genomics) according to the manufacturer’s instructions. Between 5,000 and 17,000 cells, depending on the donor, were loaded at a concentration of 700–1200 cells/µL onto a Chromium Controller instrument (10× Genomics) to generate single-cell gel bead-in-emulsions (GEMs). In this step, each cell was encapsulated with primers containing a fixed Illumina Read 1 sequence, a cell-identifying 16-bp 10× barcode, a unique molecular identifier (UMI), and a poly-dT sequence. Upon cell lysis, reverse transcription yielded full-length, barcoded cDNA, which was then released from the GEMs, amplified using polymerase chain reaction, and purified using magnetic beads (SPRIselect, Beckman Coulter). Enzymatic fragmentation and size selection were used to optimize the cDNA size prior to library construction. Fragmented cDNA was then end-repaired, A-tailed, and ligated to Illumina adaptors. A final polymerase chain reaction amplification using barcoded primers was performed for sample indexing. Library quality control and quantification were performed using a Qubit 3.0 Fluorometer (Life Technologies) and an Agilent 4200 TapeStation System (Agilent), respectively. Sequencing was performed on NextSeq500 and NextSeq2000 instruments (Illumina) at an average depth of 30,000 reads/cell.

### Single-cell RNA-seq analysis

The scRNA-seq data was demultiplexed and aligned to the human reference genome (GRCh38). The feature-barcode matrix was quantified using Cell Ranger (v6.0.1) from 10X Genomics. Computational analysis was carried out using Seurat (v4.2.0). In order to remove the possible heterotypic and homotypic doublets/multiplets, cells were analyzed with scrublets^23^ and removed. Cells underwent quality control filters based on the number of detected genes, number of unique molecular identifiers (UMIs), and the proportion of UMIs mapped to mitochondrial and ribosomal genes per cell. Each dataset was normalized, highly variable genes were identified, and unwanted sources of variation were removed. Integration of all datasets was performed using Seurat’s canonical correlation analysis. Samples from del(5q) MDS patients and healthy donors were integrated using the Seurat pipeline. Counts were log normalized and scaled, and integration was performed on the 2,000 genes with the highest variability across samples, using 50 dimensions. Nonlinear dimensionality reduction was conducted using UMAP. To characterize the cell types and states defined by each cluster, we manually reviewed the differentially expressed genes (DEGs) identified for each cell cluster using canonical marker genes as a reference.

### 5q deletion analysis

To differentiate del(5q) cells from non-del(5q) cells, CopyKat^24^ and CaSpER^25^ were applied. CaSpER was used with a non-del(5q) MDS sample that had normal karyotype^26^ as reference, whereas for CopyKat, a reference composed of a combination of three healthy samples was employed. To validate the generated results, we also analyzed a non-del(5q) MDS sample using the same reference as a negative control. The raw results from CaSpER underwent filtering by extracting large-scale events using a threshold of 0.75. Subsequently, results were binarized as described in the tool’s methods, classifying each arm of each chromosome’s arm as amplified, neutral, or deleted. For the CopyKat results, a cell clustering was performed on the results from CopyKat based on the values of copy number alterations in 220 Kbp bins of the targeted region, obtaining a cluster composed of cells with negative values. Cells exhibiting negative copy number alterations in the chr5 q15-31 region, as determined by both methods, were classified as del(5q) cells. Conversely, cells for which no alterations were identified by either method were classified as non-del(5q) cells.

### Gene Regulatory Network analysis

For each comparison, 100 transcription factors and 1,000 target genes were selected based on their variability, determined by calculating the maximum absolute deviation. To determine the optimal parameters, each analysis involved a cross-validation run of SimiC^27^. The resulting Gene Regulatory Networks (GRNs) were visualized using the GRN incidence matrices provided by SimiC. Histograms for different regulons were computed using the “regulon activity score” provided by SimiC. Additionally, this score was utilized to calculate the regulatory dissimilarity score for the selected cell clusters.

### Differential expression analysis

Different methodologies were employed depending on the specific contrasts being examined. When the contrast allowed for the comparison of cells from different samples within each phenotype, we utilized the Libra^28^ framework in combination with edgeR-LRT^29^. This approach involved generating pseudobulks for each cell type, per sample, effectively aggregating the expression profiles of cells from the same phenotype. By utilizing edgeR-LRT, statistical testing was performed to identify genes that exhibited significant differential expression while accounting for the batch effect. In cases where the contrast involved a limited number of samples in any of the phenotypes, we used MAST^30^ (Model-based Analysis of Single-cell Transcriptomics) methodology.

### Cell to cell communication analysis

The analysis of cell-to-cell communication using Liana^31^ (V0.1.8) was carried out individually for each sample, comparing the healthy samples with del(5q) MDS samples. Interactions were filtered based on their statistical significance (p-value < 0.05), using the p-value from CellChat and CellPhoneDB, and on the magnitude of the interaction (log10 > 5).

## RESULTS

### Identification of CD34**^+^** cells harboring del(5q) in MDS patients using scRNA-seq data

To identify the transcriptional alterations characterizing hematopoietic progenitors harboring del(5q), we initially performed scRNA-seq of CD34^+^ cells of four newly diagnosed patients with del(5q) MDS (Patient_1-4), and three age-matched healthy donors (Healthy_1-3) using the 10X Genomics technology. The clinical and genomic characteristics of the MDS patients and healthy donors are shown in Table S1. The percentage of cells with del(5q) based on the cytogenetic analysis varied between 35 to 90%. In all cases, the common deleted region encompassed bands 5q13-33 (genes shown in Table S2).

A total of 55,119 and 45,311 cells from patients and healthy donors, respectively, were profiled and integrated. After applying quality filters, 46,772 and 43,442 cells were eventually included in the downstream analysis (Fig. 1A). Data was integrated, clustered and manually annotated (Fig. 1B-C, Fig S1B-C) based on curated markers (Fig. 1D), obtaining 14 and 13 clusters (patients and donors, respectively) representing all the expected hematopoietic progenitor subtypes. Contribution of every MDS patient and donor to the composition of all the clusters was identified (Fig. 1E, Fig. 1SE), although each patient showed different proportions of hematopoietic progenitors (Fig. 1F). In the case of healthy donors, the same analysis was performed (Fig. S1A-E). Although there were some differences in the percentage of hematopoietic progenitors between MDS patients and healthy donors (e.g., HSC), these differences were not statistically significant which might be related to the high variability in cell composition across samples (Fig. 1G).

**Figure 1.**
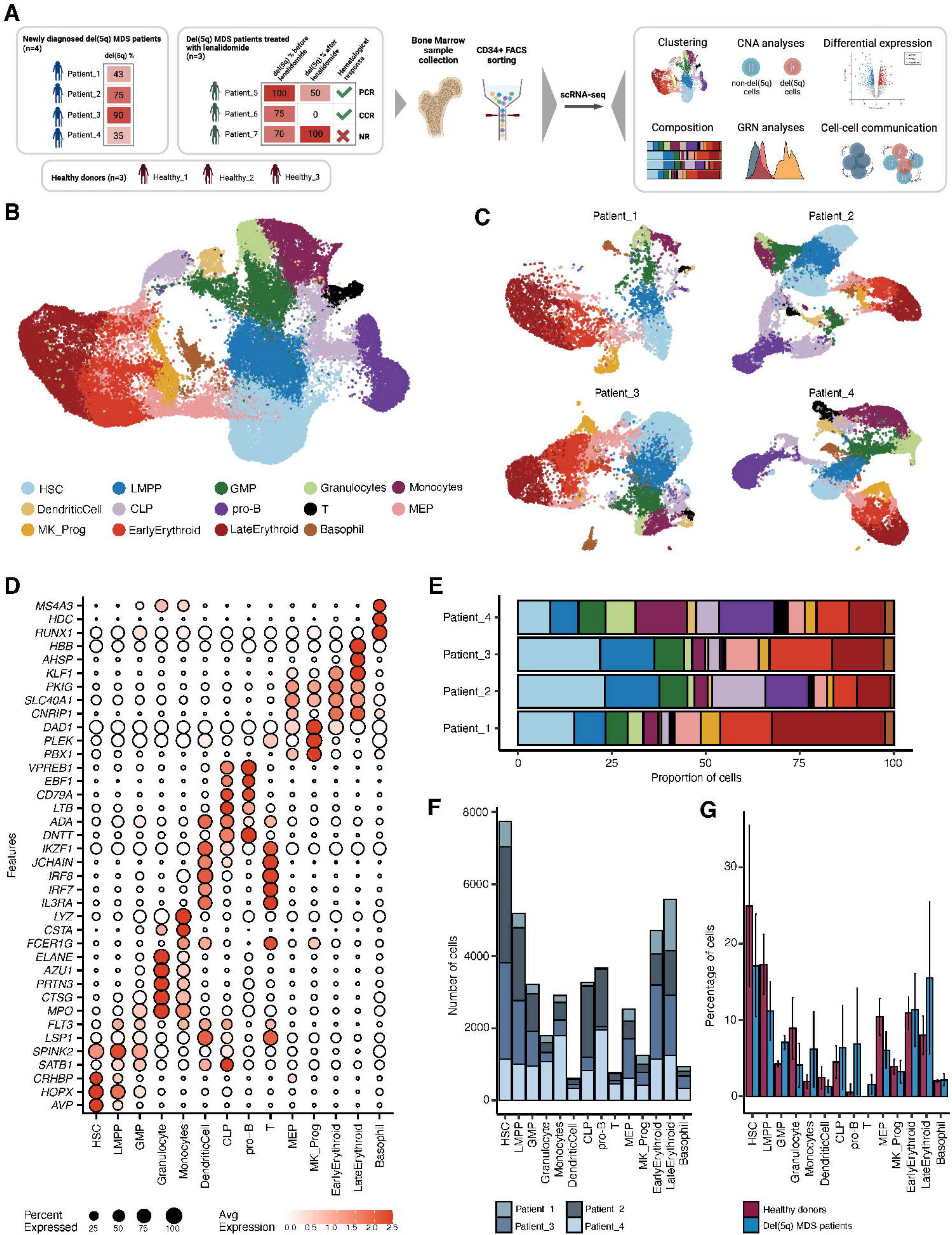
Haematopoietic CD34+ cells from four independent del(5q) MDS patients were assayed by scRNAseq. **(A)** CD34+ cells were obtained from bone marrow aspirates of newly diagnosed del(5q) MDS (n=4), healthy donors (n=3) and patients treated with Lenalidomide (n=3) and subjected to single-cell RNA sequencing and analysis. PCR: partial cytogenetic responder; CCR: complete cytogenetic responder; NR: non-responder. **(B)** An overview of the 42,494 cells that passed quality control and filtering for the subsequent analysis in this study. Uniform Manifold Approximation and Projection (UMAP) represents the 14 clusters that were analyzed. **(C)** Per patient UMAP showing the identity of the cells projected from the integrated space. **(D)** Dotplot showing the percentage and value of the normalized expression of the canonical marker genes used to assign the cell identity to each cluster. **(E)** Barplot representing the contribution of cells from each patient to the different cell types. **(F)** Barplot representing the number of cells assigned to each cell type for the studied patients. **(G)** Barplot representing the percentage of cells assigned to each cell type for the del(5q) MDS patients (n=4), and the healthy samples (n=4). No significant differences were found (pvalue<0.05 on Wilcox test) between the number of cells in the two phenotypes, except for T cells, which were not detected in healthy samples.

In order to detect individual cells harboring del(5q), we performed a copy number alteration (CNA) analysis using two different algorithms: CopyKat^24^ (Fig. 2A, see Methods), and CaSpER^25^ (Fig. 2B, see Methods). Cells that harbored del(5q) according to both methods were annotated as del(5q) cells (Fig. 2C). To validate this classification, we analyzed the expression pattern of genes encoded in the deleted region in individual cells. Due to the sparsity of scRNA-seq data, we were only able to detect expression of six genes, *CD74*, *RPS14, BTF3, COX7C, HINT1* and *RPS23* which was decreased in del(5q) when compared to non-del(5q) cells at sample level (Fig. 2D), further confirming our del(5q) cell classification. Interestingly, for each individual patient, the proportion of del(5q) in the CD34^+^ progenitor cells was consistent with that obtained by karyotype in total bone marrow (Fig. 2E).

**Figure 2.**
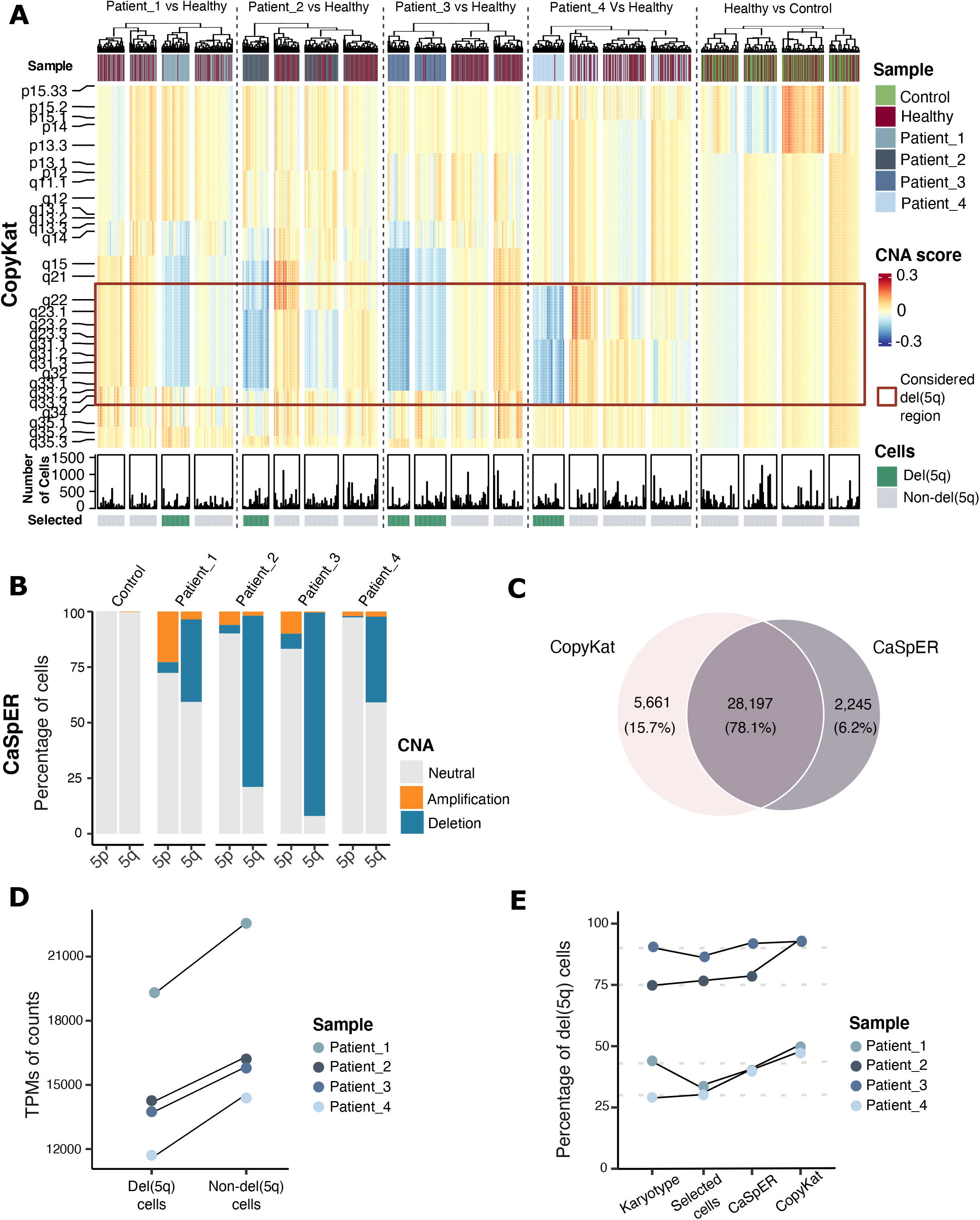
Identification of cells harboring del(5q) deletion in MDS patients. **(A)** Heatmap of the results of CopyKat showing the copy number alteration score given to each 200 kb bins in chromosome 5. In order to represent cells, a clustering has been performed within each sample (kmeans with k=80), and a posterior clustering has been applied to detect the clusters containing the cells harboring the deletion. The control sample represents a MDS sample with normal karyotype used as control for the algorithm, and the healthy sample represents an additional negative control with normal karyotype. **(B)** Barplot representing the percentage of cells for each sample presenting an amplification, a deletion, or a normal number of copy number variation, obtained using CaSpER for each branch of chromosome 5. The control corresponds to a MDS sample with normal karyotype used as a reference by the algorithm. **(C)** Venn diagram representing the numbers and percentage of cells classified as del(5q) by both algorithms. **(D)** Pseudobulk normalized expression of 6 genes (*CD74, BTF3, COX7C, HINT1, RPS23, RPS14*) showing high expression on our datasets separated by genotype. The pseudobulk values were calculated using the del(5q) and non-del(5q) cells independently. **(E)** Graph depicting the percentages of cells assigned for each sample as del(5q) by the different methods (karyotype, CaSpER, or CopyKat). Selected cells correspond to the cells detected by both CaSpER and CopyKat as those harboring a negative alteration of the number of copies on the studied region.

We then interrogated the distribution of del(5q) cells across the different hematopoietic progenitors. Cells with the deletion were detected in all the defined hematopoietic progenitor clusters (Fig. 3A), although a high heterogeneity of distribution was observed among patients (Fig. 3B-C). Despite the observed heterogeneity, a statistically significant accumulation of del(5q) cells was detected in early erythroid progenitors across all individuals (hypergeometric test, FDR < 0.05). Additionally, three out of four patients exhibited statistically significant enrichment of del(5q) in GMPs, megakaryocyte and late erythroid progenitors (Fig. 3D). Collectively, these results indicated a bias of del(5q) cells towards specific myeloid compartments, mainly towards erythroid cells, which is consistent with the association between this genetic lesion and the anemia that characterizes patients with del(5q) MDS.

**Figure 3.**
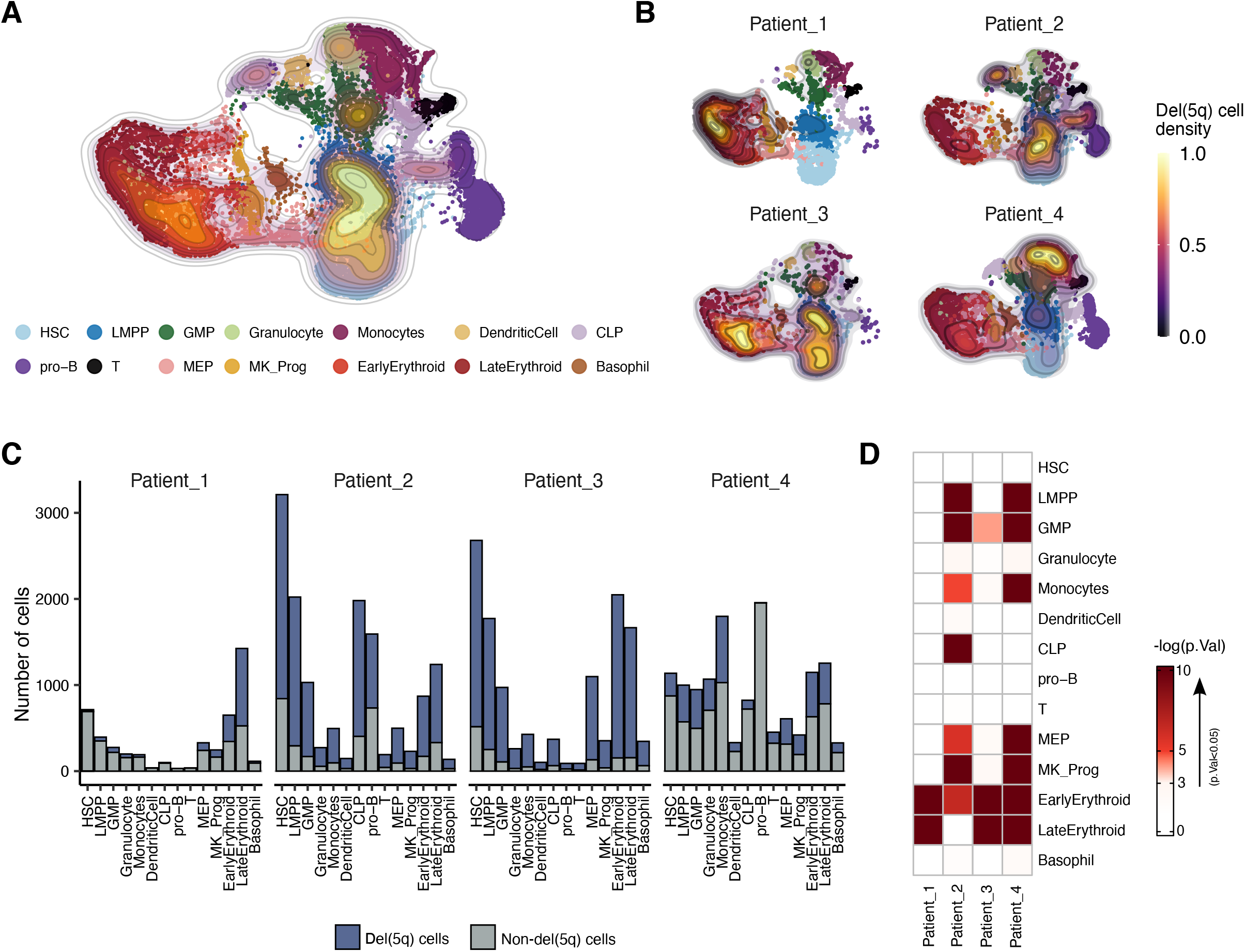
Distribution of del(5q) cells within the CD34^+^ cells of MDS patients. **(A)** UMAP representing all the MDS samples integrated and colored by sample. The density map represents the distribution of the cells classified as del(5q) by the two algorithms. **(B)** UMAPs with density maps representing the distribution of del(5q) cells per individual patient. **(C)** Barplots showing the number of del(5q) and non-del(5q) cells composing each cell type for each MDS patient. **(D)** Heatmap representing the p-values (-log(p-value)) of the hypergeometric tests used to evaluate the enrichment of del(5q) cells in each different cell type. Any color different from white represents an enrichment of del(5q) cells.

### Transcriptional differences between del(5q) and non-del(5q) cells within MDS patients are driven by few specific transcriptional programs playing key roles in MDS

To delve into the transcriptional program associated with del(5q) cells in patients with MDS, we performed a pseudobulk differential expression (DE) analysis between del(5q) and non-del(5q) cells for each cell population. Intriguingly, considering every type of hematopoietic progenitors, only seven genes in del(5q) were differentially expressed (downregulated) in comparison with non-del(5q) cells (Fig. 4A). Some of the downregulated genes played a key role in MDS and other non-hematological tumors, such as *PRSS21,* which encodes for a tumor suppressor frequently hypermethylated in cancer^32^, *MAP3K7CL*, whose downregulation serves as a biomarker in other types of cancer^33^, and *CCL5*, whose downregulation is associated with high-risk MDS^34^.

**Figure 4.**
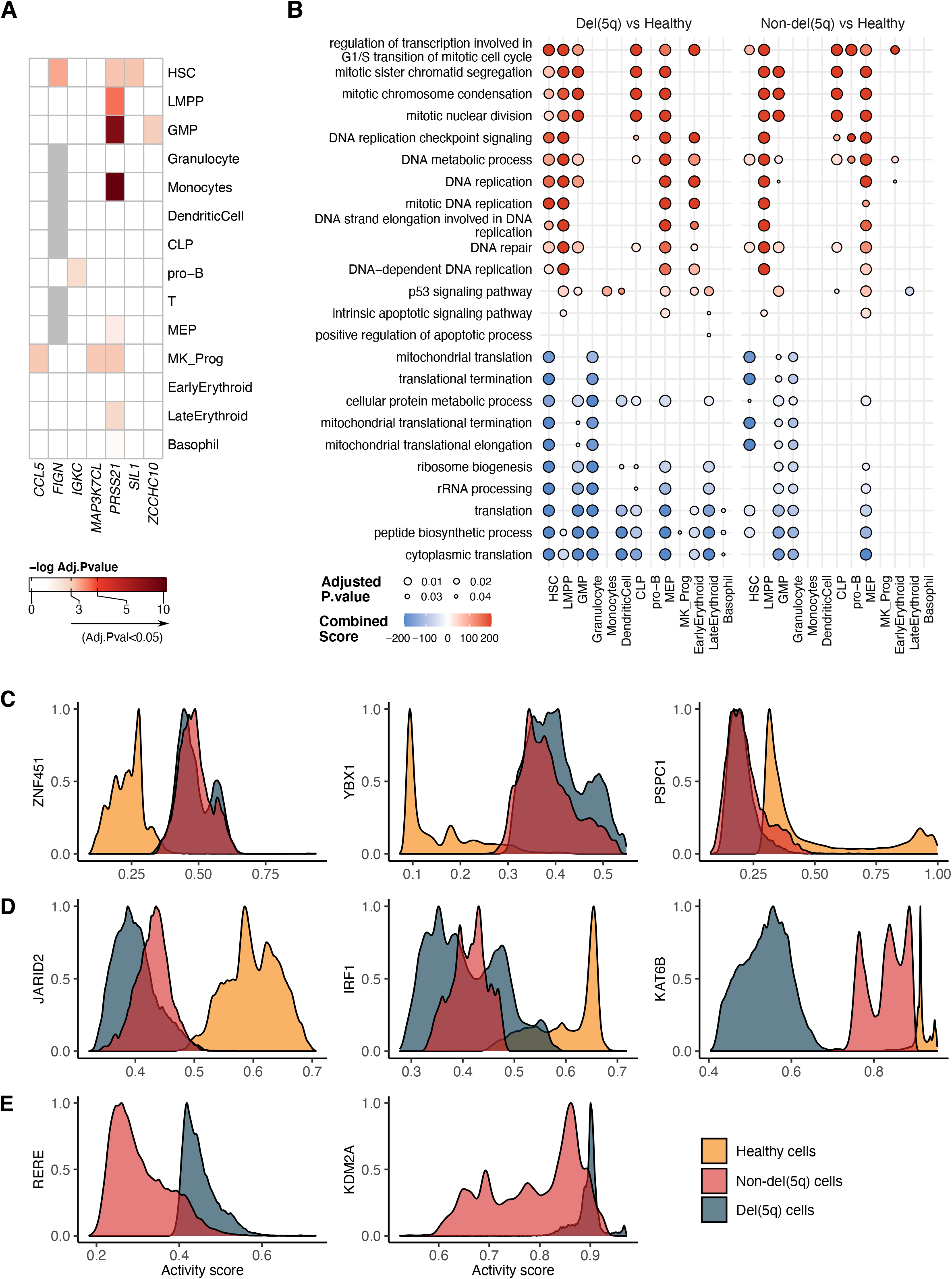
Differential expression analysis between del(5q) and non-del(5q) cells within MDS samples exposes transcriptional similarities. **(A)** Heatmap representing the differentially expressed genes (pvalue<0.05 and |logFC|>2) observed in all the samples. To generate the heatmap, the four del(5q) MDS were combined generating a pseudobulk per each cell type. **(B)** Dotplot of the over-representation analysis of the differentially expressed genes (adj. pval< 0.05) obtained for the del(5q) versus Healthy, and the non-del(5q) versus Healthy contrasts. **(C-E)** Histograms representing the activity score in all the cells separated by conditions. Some regulons behaved similarly in the MDS samples (non-del(5q) and del(5q) cells) compared to healthy cells **(B)**, while other regulons behaved differently in the three different conditions **(C).** Some inferred regulons had an activity score on the MDS samples, while lacking on the healthy samples **(E)**.

Due to the unexpected transcriptional similarity between del(5q) and non-del(5q) cells within MDS patients, we next performed a DE analysis between del(5q) MDS cells and CD34^+^ cells from healthy donors. This comparison yielded 20 to 988 differentially expressed genes (FDR<0.05 and |logFC|>2), depending on the progenitor cell (Fig. S2A, Table S3). Although most of these genes were cell-type-exclusive, they were enriched in similar pathways in most of the cell types (Fig. 4B, left panel). Genes overexpressed in del(5q) cells were enriched in cell cycle and mitosis-related signatures, such as DNA replication and mitotic nuclear division, and showed increased DNA repair, suggesting that loss of 5q confers increased proliferative potential. Additionally, del(5q) erythroid progenitors, LMPPs, GMPs, DCs, and monocyte progenitors showed a positive enrichment in p53 signaling pathway, and an enrichment in apoptosis-related pathways was detected in LMPP, MEP and late erythroid progenitors, but not in early erythroid progenitor cells. Our results are in line with the increased levels of apoptosis described for del(5q) patients^35–37^. Downregulated genes showed enrichment in processes related to ribosomes and translation in all hematopoietic progenitors, in line with previous works that have described del(5q) MDS as a ribosomopathy^2,36,38^. Interestingly, besides the cytoplasmic translation, we also observed mitochondrial translation altered in HSCs, GMPs and granulocyte progenitors. The comparison of non-del(5q) and healthy cells resulted in 64-736 genes per progenitor (Fig. S2B, Table S3). Enriched processes were also homogeneous among most hematopoietic progenitors and, as expected, were similar to the ones observed in del(5q) vs healthy (Fig. 4B, right panel).

Despite the low number of DE genes between del(5q) and non-del(5q) cells, we were interested in understanding whether differences in GRN might be observed between these two populations. Unlike DE analysis, which is performed in a gene-by-gene manner, GRN studies use data-driven grouping of genes to enable the identification of mechanistic transcriptional differences between conditions. Thus, we applied SimiC^27^ to compute the regulatory activity of regulons and observed that although some regulons behaved uniformly (low regulatory dissimilarity score, in Fig. S3A black-purple color) between the three conditions (del(5q), non-del(5q) and healthy cells), a group of regulons showed differential activity (high regulatory dissimilarity score, in Fig. S3A yellow-orange color) across the conditions (Fig. S3A). Among them, three different regulon activity patterns arose. Firstly, a group of regulons that showed similar activity between non-del(5q) and del(5q) cells, and different to healthy cells, in line with DE analyses, such as the ones driven by *ZNF451*, *YBX1* and *PSPC1* (Fig. 4C). Secondly, there were regulons with differential activity between the three conditions, such as those driven by *JARID2*, *IRF1* and *KAT6B*, among others (Fig. 4D). The three regulons showed high activity in healthy elderly cells, whereas they presented a progressively lower activity in non-del(5q) cells, and their lowest activity in del(5q) cells. *JARID2* acts as a tumor suppressor and plays a crucial role in the leukemic transformation of myeloid neoplasms^39^, and its deletion promotes an ineffective hematopoietic differentiation^40^, suggesting that the low activity of this regulon may negatively impact the hematopoietic differentiation of these patients. *IRF1* is located in 5q31.1 and its deletion in one or both alleles have been observed in MDS and AML patients with chromosome 5 abnormalities^41^. *IRF1* has been described as a master HSC regulator, and its loss impairs HSC self-renewal and increases stress-induced cell cycle activation, suggesting that its low activity in patients could confer proliferative advantage^42^. Decreased expression of *KAT6B* in aged hematopoietic stem cells has been associated with impaired myeloid differentiation^43^, suggesting that its almost non-existent activity in del(5q) cells may contribute to aberrant differentiation of these cells. Lastly, we detected regulons exhibiting differential activity between del(5q) and non-del(5q) cells and that showed no activity in healthy cells. In particular, regulons driven by *RERE* and *KDM2A* showed higher activity in del(5q) cells than in non-del(5q) cells (Fig. 4E). *RERE* negatively regulates the expression of target genes, and such genes were enriched in cytoplasmic translation, ribosome biogenesis and ribonucleoprotein complex biogenesis pathways, among others (Fig. S3B). The *KDM2A* regulon was enriched in protein stabilization, regulation of cellular protein catabolic process and regulation of protein stability (Fig. S3B). The association of *KDM2A* and ribosomal genes has been already described by previous studies, postulating that *KDM2A* overexpression reduces the transcription of rRNA^44,45^.

Altogether, our results suggest a low transcriptional impact of 5q loss, with del(5q) and non-del(5q) cells presenting very similar gene expression alterations when compared to healthy controls, with such alterations being involved in processes that could contribute to abnormal hematopoietic differentiation. Nevertheless, although limited in number, genes and regulons specifically altered in del(5q) cells, such as those driven by *JARID2*, *KAT6B*, *RERE* or *KDM2A*, seem to be relevant for proliferation and myeloid differentiation, supporting the concept that cells harboring the deletion may have a more prominent role in the promotion of altered hematopoiesis.

### Abnormal cell-to-cell communication in del(5q) progenitors

To investigate whether the 5q deletion has a detrimental effect on cell-cell interactions between CD34^+^ progenitors, thus contributing to disease development, we performed a cell-to-cell communication analysis using Liana^31^ in both del(5q) and healthy controls datasets. We identified 4,534 interactions in healthy controls, and 314 interactions that were common to all del(5q) MDS patients, most of them overlapping with those found in healthy cells (Fig. 5A). Despite this strong overlap, several differences between del(5q) MDS and healthy individuals were detected: in patients, monocyte progenitors were the most communicative cells, interacting mainly with early erythroid progenitors (Fig. 5B). However, in healthy donors, HSCs, GMPs, DC, monocyte and granulocyte progenitors were the most interactive compartments, with a notable communicative pattern between granulocyte and GMP/DC progenitors (Fig. 5C). Furthermore, genes involved in these differential interactions were overrepresented in different biological processes in each phenotype. For instance, interactions driven by healthy hematopoietic progenitors were enriched in negative regulation of apoptosis, HSC proliferation, leukocyte/DC differentiation, and hemopoiesis, whereas those found in MDS progenitors were enriched in negative regulation of translation, oncogenic MAPK signaling and HIF-1 signaling (Fig. 5D). Focusing on interactions driven by del(5q) and non-del(5q) cells within the patients (Fig. 5B), we observed very subtle differences regarding the communicational pattern and the number of interactions observed for each of the compartments, and there were no interactions specifically established between del(5q) cells, corroborating the high similarity already described between del(5q) and non-del(5q) cells. Overall, our results are consistent with the previously described lack of significant differences in gene expression between del(5q) and non-del(5q) cells, suggesting that deregulation of hematopoiesis in patients with 5q MDS affects all CD34^+^ cells.

**Figure 5.**
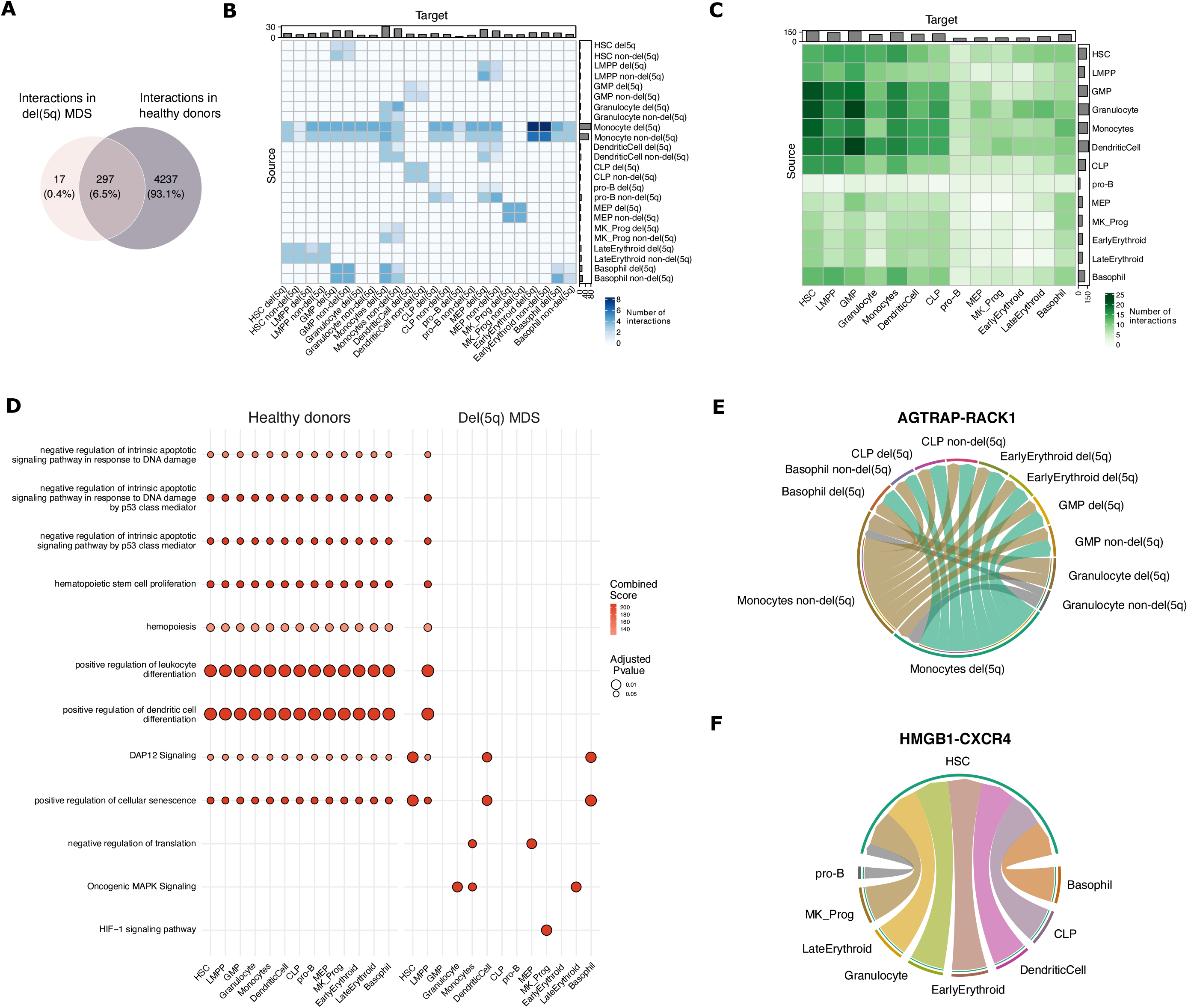
Cell to cell communication reveals shared and unique interactions between the three conditions. **(A)** Venn diagram showing the number of unique interactions in MDS and healthy samples. Healthy unique interactions are all the interactions present in at least one of the healthy samples, while MDS unique interactions are the interactions present in all the patients. **(B-C)** Interactions for each cell type in healthy **(B)** and MDS samples **(C)** were quantified using Liana, taking into account the direction (source-target), and were represented in a heatmap. **(D)** Dotplot representing the pathways to which belong the genes mediating the communication in the healthy samples, and the genes in the exclusive communications in the MDS samples. The color represents the combined score and the size of the dot indicates the adjusted pvalue. **(E-F)** Chord diagrams representing specific communications between different cell types, where the inner circle represents the target cell type and the outer circle represents the source cell type. The interactions represented are for the AGTRAP-RACK1 pair in the MDS samples **(E)** and HMGB1-CXCR4 pair in the healthy samples **(F)**.

To uncover specific interactions that may contribute to the disease, we next focused on those interactions that had been gained or lost in MDS versus controls. There were 17 novel interactions identified in patients that were totally absent in healthy individuals, suggesting that additional communications arise when developing the disease. For instance, *AGTRAP* expressed in monocyte and late erythroid progenitors interacted with *RACK1* in HSCs, LMPPs, MEPs, pro-B and basophil progenitors (Fig. 5E). *AGTRAP* is known to be implicated in hematopoietic cell proliferation and survival^46^, whereas *RACK1* has been postulated as a potential therapeutic target for promoting proliferation in other myeloid neoplasms^47,48^. The fact that these molecules are highly expressed in MDS could potentially be contributing to the enhanced proliferation observed in MDS cells. In contrast, there were 37 interactions that appeared in the healthy donors and were absent in the patients, including the one established between *HMGB1* expressed in CLPs, DC, granulocyte, basophil, megakaryocyte, early erythroid and late erythroid progenitors, and *CXCR4* present in HSCs (Fig. 5F). *HMGB1*-*CXCR4* interaction is known to trigger the recruitment and activation of inflammatory cells in tissue regeneration^49,50^, thus its loss could have a negative impact on the bone marrow niche. In summary, these analyses may allow the identification of potential interactions implicated in the pathogenesis of the disease that could represent new therapeutic targets.

### Effect of Lenalidomide treatment on the transcriptional programs of del(5q) and non-del(5q) cells from MDS patients

We next aimed to understand the effect of treatment with the standard-of-care, lenalidomide, on the transcriptional alterations observed in del(5q) and non-del(5q) cells. We performed scRNA-seq on CD34^+^ cells of two patients (Patient_5-6), which had achieved hematological response (one with partial cytogenetic response (PCR), and the other one with complete cytogenetic response (CCR), respectively) (clinical information in Table S1). Data were integrated clustered, manually annotated, and del(5q) cells were identified as described before (Fig. 6A). Patients showed different percentages of del(5q) cells which were consistent with karyotype results (Fig. 6C): the patient with PCR showed 1,939 cells with del(5q) (37.13%), whereas the patient showing CCR presented only 11 cells with del(5q) after treatment (0.15%), validating the persistence of del(5q) progenitor cells at the time of complete clinical and cytogenetic remission^9^. Similar to what we observed at diagnosis, the distribution of del(5q) cells was heterogeneous among patients (Fig. 6A-B), and both responders exhibited a statistically significant del(5q) enrichment in GMPs and erythroid progenitors. Interestingly, patient with PCR also showed an enrichment in LMPPs, megakaryocyte, monocyte, and granulocyte progenitors (Fig. 6D).

**Figure 6.**
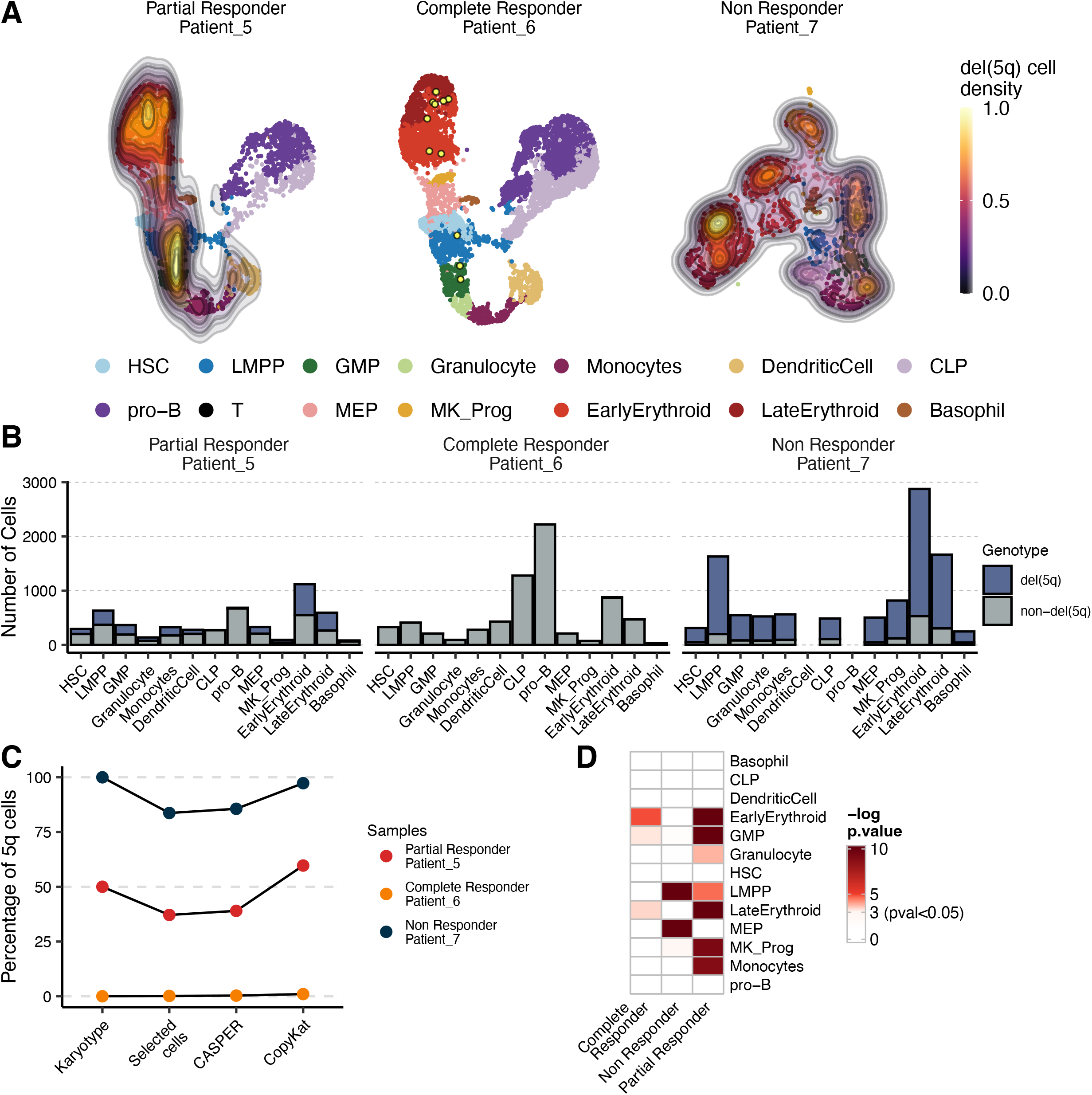
Distribution of del(5q) cells on the CD34^+^ cells after lenalidomide treatment. **(A)** UMAP colored by cell type and with the density lines and colors representing the distribution of del(5q) cells (selected by CASPER and CopyKat) in the different post-treatment samples. **(B)** Barplot showing the number of cells per cell-type, colored by the presence or absence of the deletion of the 5q region and separated by sample. **(C)** Percentage of the cells identified as del(5q) by karyotype, CASPER, CopyKat, and the selection by intersecting the two methods. **(D)** Heatmap representing the p-values (-log(p-value)) of the hypergeometric tests used to evaluate the enrichment of del(5q) cells in each different cell type for the three post-treatment samples. Any color different from white represents an enrichment of del(5q) cells.

We have demonstrated in the previous analyses that at diagnosis both del(5q) and non-del(5q) progenitors displayed transcriptional profiles linked to an aberrant hematopoiesis. Since both PCR and CCR patients were in hematological response, we hypothesized that the remaining CD34^+^ cells after lenalidomide treatment, which are mainly composed of non-del(5q) progenitors, must be able to promote improved hematopoiesis and thus restore the transcriptional profile of normal progenitor cells. To demonstrate that lenalidomide, besides the potential apoptosis of del(5q) cells, could reverse transcriptional alterations harbored by non-del(5q) cells in responder patients, we performed a DE analysis between non-del(5q) cells from the CCR and PCR and the four patients at diagnosis at pseudobulk levels This comparison revealed significant transcriptional changes after lenalidomide treatment, resulting in 622-3,609 genes in the different progenitor populations for the CCR (Fig. S2C, Table S3) and between 458-2,409 genes (FDR<0.05 and |logFC|>2) for the PCR (Fig. S2D, Table S3). Note that these transcriptional differences are significantly greater than those related to patient heterogeneity at diagnosis (see previous sections), indicating that most uncovered altered genes after treatment are probably due to treatment effect rather than patient heterogeneity. Genes altered upon treatment were enriched in ubiquitination and proteasome-mediated catabolic processes, and in phosphatidylinositol related pathways, which is in line with the mechanism of action described for lenalidomide in (5q) MDS patients^51,52^. Moreover, we detected an enrichment in autophagy-related processes. Overall, our results suggested an increase of the two most important protein degradation pathways in non-del(5q) cells upon lenalidomide treatment (Fig. 7A, first and second panels, Table S4). Furthermore, hematopoietic progenitors exhibited enhanced erythropoietin signaling and an increased expression of genes involved in erythroid differentiation after treatment (Fig. 7B), validating the enhanced erythropoiesis in response to treatment^19^. Our analyses also detected a positive enrichment of *PD-L1* expression and *PD-1* checkpoint in non-del(5q) cells of the responder patients after treatment, suggesting a potential immunosuppressive mechanism of these cells in response to lenalidomide (Fig. 7A, first and second panels, Table S4).

**Figure 7.**
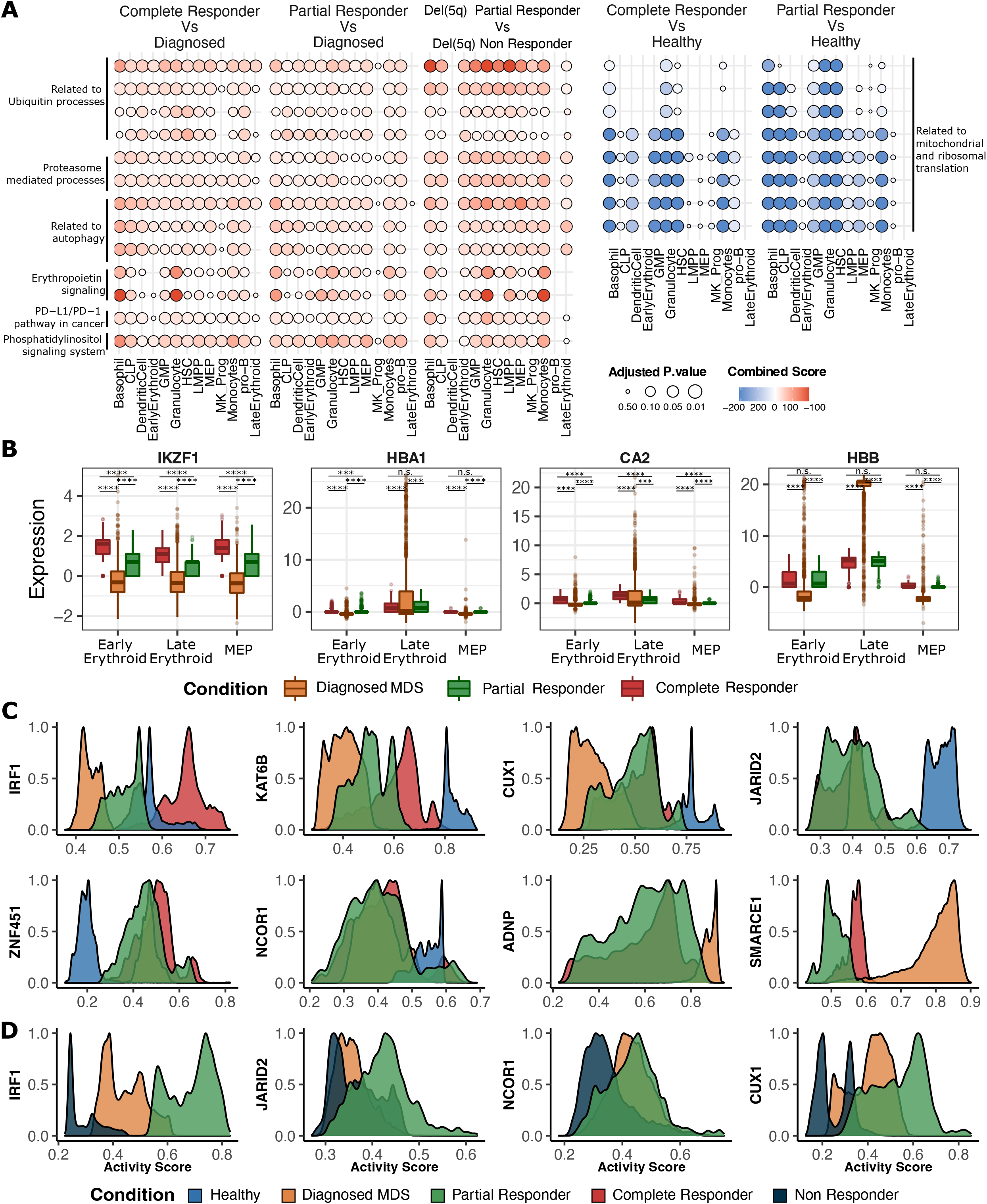
Differential expression between treated and untreated patients unravels persistent transcriptional alterations after lenalidomide treatment. **(A)** Dotplot showing an over representation analysis performed with the differentially expressed genes (pval<0.05 and avgLogFC>2) obtained in different comparisons: non-del(5q) cells of the complete responder vs at diagnosis (1st panel); non-del(5q) cells of the partial responder vs at diagnosis (2nd panel); del(5q) cells of the partial responder vs the non-responder (3rd panel); non-del(5q) cells of the complete responder vs healthy cells (4th panel); non-del(5q) cells of the partial responder vs healthy cells. (B) Boxplot showing the normalized expression for non-del(5q) cells of erythroid differentiation-related genes in the del(5q) MDS samples at diagnosis or after treatment with lenalidomide (partial and complete responder). The significance level has been calculated with Wilcoxon (n.s.: p > 0.05; *: p ≤ 0.05, **: p ≤ 0.01, ***: p ≤ 0.001, ****: p ≤ 0.0001). (C) Activity scores calculated on the non-del(5q) cells for different transcription factors associated with differentiation for the different samples: healthy samples, del(5q) MDS at diagnosis and after treatment with the lenalidomide. **(D)** Activity scores calculated on the del(5q) cells for different GRNs associated with differentiation in del(5q) cells of patients at diagnosis, with a partial response and a non-responder after lenalidomide treatment.

GRN analyses evidenced that some of the alterations described at diagnosis were reverted after treatment in non-del(5q) cells. *IRF1*, the master HSC regulator located in 5q31.1, which showed abnormally low activity at diagnosis in non-del(5q) cells, showed an increased activity after treatment in both patients, with the PCR not reaching the activity level seen for healthy cells, and the CCR showing an augmented activity comparable to the healthy cells (Fig. 7C). *KAT6B*, whose lower expression has been associated with impaired myeloid differentiation, showed an augmented activity in both patients despite not reaching the activity level of healthy cells. Finally, *CUX1*, a TF frequently mutated in myeloid malignancies and whose knockdown leads to an MDS-like phenotype^53^, presented similar activity in non-del(5q) cells, showing higher activity than at diagnosis (Fig. 7C).

Importantly, although some transcriptional lesions were reverted upon lenalidomide treatment, non-del(5q) cells continue exhibiting ribosome-related alterations, showing a negative enrichment of processes related to ribosomes, translation, and mitochondrial translation when compared to healthy cells (Fig. 7A, fourth and fifth panels, Fig. S2E-F, Table S3-4). After treatment, early and late erythroid non-del(5q) progenitors from responding patients showed no statistically significant changes in these pathways. Moreover, GRN analyses detected groups of regulons with similar activity for non-del(5q) cells at diagnosis and after treatment response, but with a different activity to the healthy cells, indicating that lenalidomide did not affect their aberrant activity. Some examples included the tumor suppressor *JARID2*^39,40^, *ZNF451*, a TF whose high expression in leukemic cells has been associated with poor outcome^54^, and *NCOR1*, a regulator of erythroid differentiation^55^ (Fig. 7C). Moreover, non-del(5q) cells exhibited, both at diagnosis and after treatment, abnormal high activity of two regulons that were not active in healthy cells: *ADNP* and *SMARCE1* (Fig. 7C). Globally, these results indicate that treatment with lenalidomide has the potential to revert some of the transcriptional alterations present at diagnosis in non-del(5q) cells at least in patients that responded to lenalidomide. Nevertheless, some of the transcriptional alterations present at diagnosis were not modified which could be relevant for abnormal hematopoiesis, and potentially, for the future relapse of the patients.

In line with what has been observed in non-del(5q) cells from the PCR, the remaining del(5q) cells generally exhibited higher ubiquitination, phosphatidylinositol signaling, autophagy, and apoptosis when compared to del(5q) cells at diagnosis (Fig. S2G, Fig. S4A, Table S3,5), which is consistent with the mechanism of action of lenalidomide^51,52^. However, these cells showed reduced ribosomal and mitochondrial translation compared to diagnosis, along with diminished DNA repair capacity (Fig. S4A). This suggests that lenalidomide has not been able to fully reverse key transcriptional alterations that may underlie the ribosomopathy characterizing the disease.

### Lenalidomide does not correct transcriptional alterations of del(5q) cells of a refractory MDS patient

Finally, to understand the transcriptional alterations associated with a lack of hematological response after lenalidomide treatment, we performed scRNAseq on CD34^+^ cells of an additional patient (Patient_7), who was refractory to lenalidomide (non-responder, NR). Data were processed as described previously (clinical information in Table S1), showing 83.8% of 5q cells (Fig. 6A-C), and a statistically significant increase of del(5q) cells in LMPPs, MEPs and megakaryocyte progenitors (Fig. 6D). We then analyzed the transcriptional differences between the remaining del(5q) cells of the responder that presented PCR, and those of the NR patient. This analysis identified 116-2,244 differentially expressed genes (FDR<0.05) per progenitor (Fig. S2H, Table S3). Del(5q) cells from the patient in PCR showed statistically significant enrichment in processes and pathways related to protein ubiquitination, proteasomal protein catabolic process, phosphatidylinositol and autophagosome when compared to the NR. Moreover, these cells also exhibited an increased erythropoietin signaling as a result of lenalidomide treatment when compared to the NR cells (Fig. 7A, third panel). Interestingly, these processes are similar to the ones detected for non-del(5q) cells when comparing these cells to those at diagnosis (see previous section), and have been described as a lenalidomide response in non-del(5q) MDS patients ^51,52^. The remaining del(5q) cells from the patient in PCR also exhibited enrichment of *PD-L1* expression and *PD-1* checkpoint pathway when compared to the refractory patient. These analyses suggested low transcriptional alterations promoted by lenalidomide treatment in the NR patient. Accordingly, DE analysis of del(5q) cells at diagnosis and after treatment in the NR patient yielded 20-121 differentially expressed genes per hematopoietic progenitor (Fig. S2I, Table S3). These few differences resulted in subtle changes in protein ubiquitination and cell cycle-related processes after treatment (Fig. S4B, Table S5), showcasing that lenalidomide did not have a high transcriptional impact on del(5q) cells of the NR.

GRN analysis demonstrated a large number of regulons that showed changes in activity after treatment in del(5q) cells from the patient in the PCR but not in the refractory patient. For example, regulons driven by *IRF1*, *JARID2*, *NCOR1*, and *CUX1*, which showed aberrant low activity at diagnosis that was partially recovered upon treatment, presented very reduced activity in the NR patient, which was lower than that observed in the PCR, and at diagnosis (Fig. 7D). Collectively, these results suggest that in NR patients lenalidomide treatment is not able to reverse part of the transcriptional lesions carried by (5q) cells, which seems to be associated with the lack of hematological response.

## DISCUSSION

Establishing the relationship between genomic and transcriptional abnormalities in hematological malignancies has allowed researchers to characterize the molecular pathogenesis of diseases such as MDS and to identify new potential targets^10–14,18^. However, for the most part, these studies have been performed using bulk sequencing data, which precludes a direct association on a per cell bases between genomic and transcriptomic alterations. Using scRNAseq data from CD34^+^ cells from patients with del(5q) MDS, we have been able to identify cells with del(5q) and non-del(5q) which enabled us to compare the transcriptional and GRN of both populations from the same patients and thus made strides in the understanding of the molecular pathogenesis of the disease.

The stem-cell origin of del(5q) MDS has been previously demonstrated^9,56^ as well as the presence of del(5q) cells even at the stage of cytogenetic response^9^. By being able to dissect the presence of del(5q) in individual cells and identifying the different type of progenitors, our study suggests uneven distribution of del(5q) in different progenitor cell populations, and also an heterogeneous distribution according to the patients. Although we cannot establish a correlation between the phenotype of the disease, the clinical symptoms, and the specific distribution of del(5q), we were able to identify an enrichment of del(5q) cells in GMP and megakaryocyte-erythroid progenitors in patients, not described to date. Future studies with larger cohorts of cases may uncover the nature of the detected enrichment and provide possible associations between the distribution of del(5q) cells and clinical characteristics of the patients.

The comparison between the transcriptional profiles between del(5q) cells and non-del(5q) within the same patient provides new insights regarding the impact of 5q deletions on pathogenesis of del(5q) MDS: 1) unlike what has been previously described in other cancers where CNAs are known to exert expression differences beyond the deleted or amplified regions, influencing genes located elsewhere in the genome^57^, the deletion did not have a high impact in the expression of other genes beyond the CDR, as only 7 differentially expressed genes were identified between del(5q) and non-del(5q) cells. 2) Interestingly, both types of cells presented similar transcriptional lesions when compared to progenitor cells from healthy individuals, indicating that in del(5q) MDS patients, both types of cells are implicated in the disease and contribute to the promotion of aberrant hematopoietic differentiation. Specifically, del(5q) and non-del(5q) cells presented alterations in biological processes described in del(5q) MDS, including cell proliferation^58,59^, p53 signaling pathway^12^ and apoptosis. In this regard, previous murine models generated by the inactivation of *RPS14* described a p53 dependent apoptosis in the erythroid lineage between basophilic/early chromatophilic erythroblasts to poly/orthochromatophilic erythroblasts^32^, but our data suggests that it could already occur at the CD34^+^ progenitor level. Furthermore, in addition to the negative enrichment in ribosome and cytoplasmic translation processes identified in our data, which are consistent with previous studies^2,36,38^, we observed an altered function of mitochondrial ribosomes in HSCs and granulocyte-lineage progenitors. Mitochondrial translation and mitoribosomes fulfill a pivotal function in cell cycle regulation and apoptosis signaling^60^, and several works have found gene expression alterations in mitoribosome or associated proteins in relation to different cancers^61^. Hence, the altered mitochondrial translation in del(5q) cells could be contributing to the molecular pathogenesis of del(5q) MDS. 3) Despite these findings, a deeper GRN analysis led to the characterization of a number of transcriptional differences between del(5q) and non-del(5q) cells, identifying a group of regulons with different activity between both genotypes. Some of the GRNs specifically altered in del(5q) cells, such as those driven by *JARID2*, *KAT6B*, *RERE* or *KDM2A*, seem to be relevant for proliferation and myeloid differentiation, supporting the concept that cells harboring the deletion may have a more prominent role in the promotion of altered hematopoiesis. Altogether, our data manifests a general transcriptional dysregulation in all the progenitor cells of del(5q) MDS patients, and shows that progenitors harboring the del(5q) deletion show additional alterations.

Lenalidomide represents the standard-of-care for transfusion-dependent del(5q) MDS patients. Our analyses suggest a close relationship between lenalidomide-mediated reversal of the transcriptional alterations harbored by hematopoietic progenitors and hematological response. In this sense, we observed that lenalidomide treatment alters the transcriptional profile of both del(5q) and non-del(5q) cells of responder patients, reverting part of the alterations observed in these cells. For example, the remaining non-del(5q) cells of responder patients show increased levels of proteasome-mediated catabolic and autophagia related processes after treatment, suggesting a compensatory increase of the two main mechanisms of intracellular protein degradation as a result of the treatment. Moreover, we detected a direct effect of lenalidomide in erythropoietin signaling, promoting the upregulation of genes involved in erythropoiesis, a mechanism that so far has been described for non-del(5q) MDS^63^. Nevertheless, certain alterations that were initially present at the time of diagnosis in del(5q) and non-del(5q) cells remain detectable at this stage of clinical evaluation, which could explain the relapses seen after the treatment. This association between reversal of transcriptional alterations and clinical response is further supported by the fact that both del(5q) and non-del(5q) cells of a non-responder patient presented very few transcriptional alterations upon lenalidomide treatment. Altogether, these results may be at odds with previous data suggesting that lenalidomide induces synthetic lethality to suppress the malignant clone without significant effect on the growth of cytogenetically normal CD34^+^ cells^11^. Our data demonstrate that lenalidomide affects both del(5q) and non-del(5q) cells, and suggest that the hematological response of del(5q) MDS patients may be due to the correction of transcriptional alterations in both types of cells.

Finally, the characterization of the transcriptional profile of remaining cells after lenalidomide treatment could help to unveil novel therapeutic strategies that could be effective in the eradication of malignant cells. In this sense, our results showed an enrichment in *PD-1/PDL-1* pathway after the treatment in non-del(5q) cells which was not detected in the NR patient, suggesting a putative immune evasion mechanism of these cells. Interestingly, lenalidomide has shown increased cytotoxic activity in combination with immune-checkpoint blockade in multiple myeloma^64^. Thus, future studies will confirm if, as our results suggest, this combination could serve as a promising therapeutic strategy for these patients.

## ACKNOWLEDGEMENTS

This work was supported by the Instituto de Salud Carlos III and co-financed by ERDF A way of making Europe (PI20/01308, PI23/00516, PI19/00726, PI22/01044), CIBERONC (CB16/12/00489), RICORS TERAV (RD21/0017/0009) and ERA-PerMed JTC 2019 (MEET-AML). MICCIN/AEI/10.13039/501100011033 and [RTI2018-101708-A-I00]. Gobierno de Navarra (AGATA 0011-1411-2020-000010/0011-1411-2020-000011). Fundación La Caixa (GR-NET NORMAL-HIT HR20-00871). Cancer Research UK [C355/A26819], FC AECC and AIRC under the Accelerator Award Program. AECC award from the Fundación AECC (INVES19059EZPO) (T.E). H2020 Marie S. Curie IF Action, European Commission, Grant Agreement No. 898356, a Ramon y Cajal contract RYC2021-033127-I MCIN/AEI/10.13039/501100011033 (M.H). Partially funded by a Ramon y Cajal contract RYC2019-028578-I, a Gipuzkoa Fellows grant 2022-FELL-000003-01, and grant PID2021-126718OA-I00 funded by MCIN/AEI/10.13039/501100011033 (I.O.) Supported by PhD fellowships from Gobierno de Navarra (0011-0537-2019-000001) (N.B.), and (0011-0537-2020-000022) (A.D.-M.); a PhD fellowship from Ministerio de Ciencia, Innovación y Universidades (FPU18/05488). We particularly acknowledge the patients and healthy donors for their participation in this study, and the Biobank of the University of Navarra for its collaboration.

## AUTHORSHIP CONTRIBUTIONS

G.S., N.B., A.D., I.O., F.P., T.E., and M.H. conceived and designed the research studies; G.S., and A.D. performed *in silico* analysis of transcriptomic data; G.S., N.B., and A.D. performed statistical analysis; G.S., N.B., A.D., P.GO. S.H-D., A.A-P., A.UA., B.A., and M.A. analyzed and interpreted data; A.VZ., P.S-M., P.A-R., J.L-E., P.A., O.C., T.J.,, A.M.,, J.M., M.D-C., D.V., and F.S. provided clinical samples and data; A.VZ., P.S-M., A.U-A., B.A., M.A. provided technical assistance; S.H-D., A.AP. and O.C. provided clinical advice; T.E., I.O., M.H., and F.P. were responsible for research supervision, coordination, and strategy. N.B., A.D. G.S., P.GO., I.O., F.P., T.E., and M.H. wrote the manuscript. All authors reviewed and approved the final version of the manuscript.

## Data and materials availability

All data needed to evaluate the conclusions in the paper are present in the paper and/or the Supplementary Materials. The scRNA-seq data generated in this study have been deposited in the Gene Expression Omnibus database (GSE245452).

