## Supplemental material for "Single cell profiling of del(5q) MDS unveils its transcriptional landscape and the impact of lenalidomide"

#### TABLES

**Table S1. Clinical information of the del(5q) MDS patients.**

**Table S2. Genomic Features of Chromosome 5 (q13.1-q33.3) Gene Loci.**

**Table S3. Differentially expressed genes per hematopoietic progenitor for several comparisons.**

**Table S4. Comprehensive breakdown of GO term categorizations from Fig 7.**

**Table S5. Comprehensive breakdown of GO term categorizations for various contrasts from Fig S4.**

Figure S1

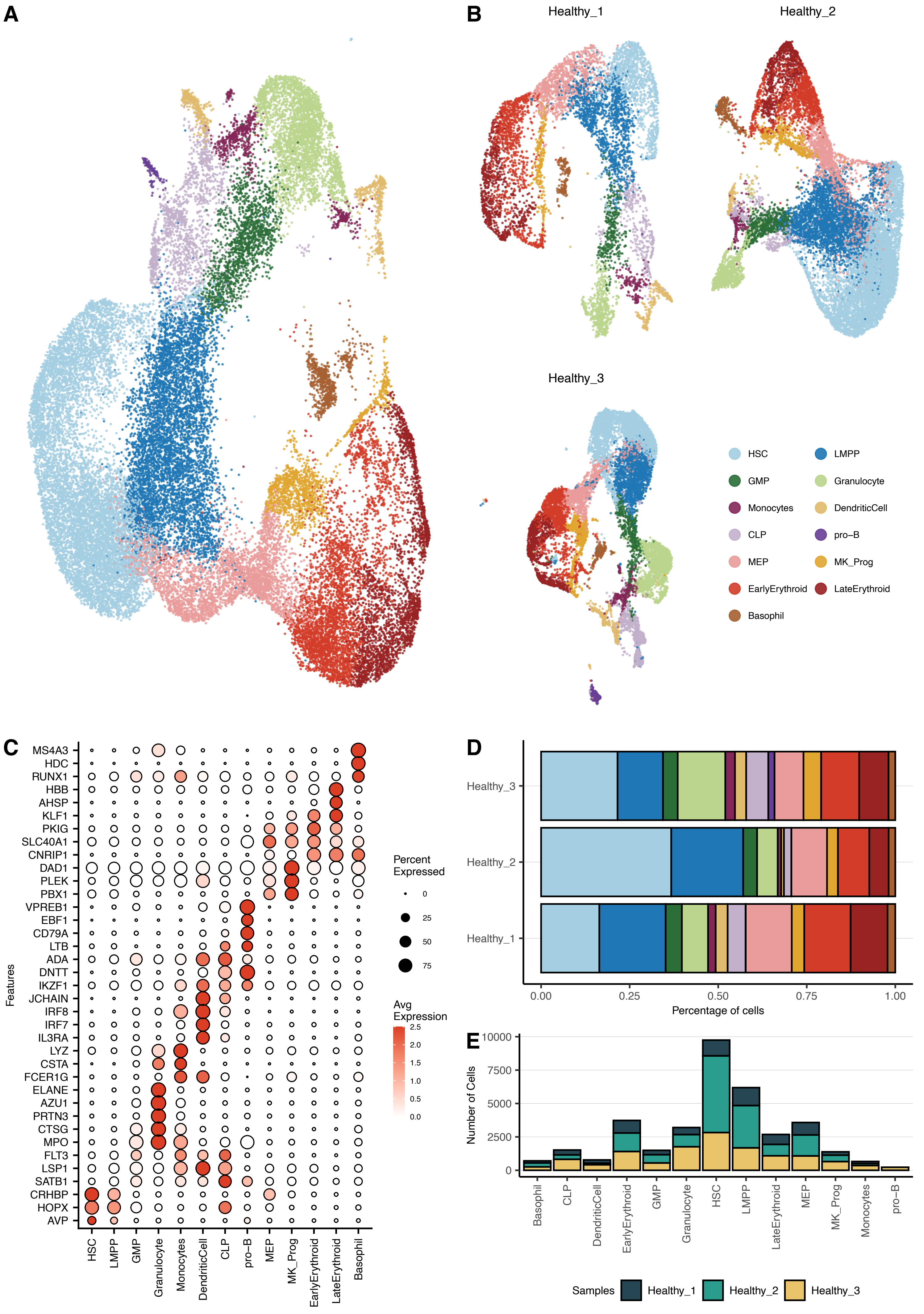
A
B
Healthy_1
Healthy_2
Healthy_3
HSC
LMPP
GMP
Granulocyte
Monocytes
DendriticCell
CLP
pro−B
MEP
MK_Prog
EarlyErythroid
LateErythroid
Basophil
C
MS4A3
HDC
RUNX1
HBB
AHSP
KLF1
PKIG
SLC40A1
CNRIP1
DAD1
PLEK
PBX1
VPREB1
EBF1
CD79A
LTB
ADA
DNTT
IKZF1
JCHAIN
IRF8
IRF7
IL3RA
LYZ
CSTA
FCER1G
ELANE
AZU1
PRTN3
CTSG
MPO
FLT3
LSP1
SATB1
CRHBP
HOPX
AVP
Features
Percent Expressed
0
25
50
75
Avg Expression
2.5
2.0
1.5
1.0
0.5
0.0
D
Percentage of cells
0.00
0.25
0.50
0.75
1.00
E
Number of Cells
10000
7500
5000
2500
0
Samples
Healthy_1
Healthy_2
Healthy_3

### Figure S2

Supplemental Figure 2

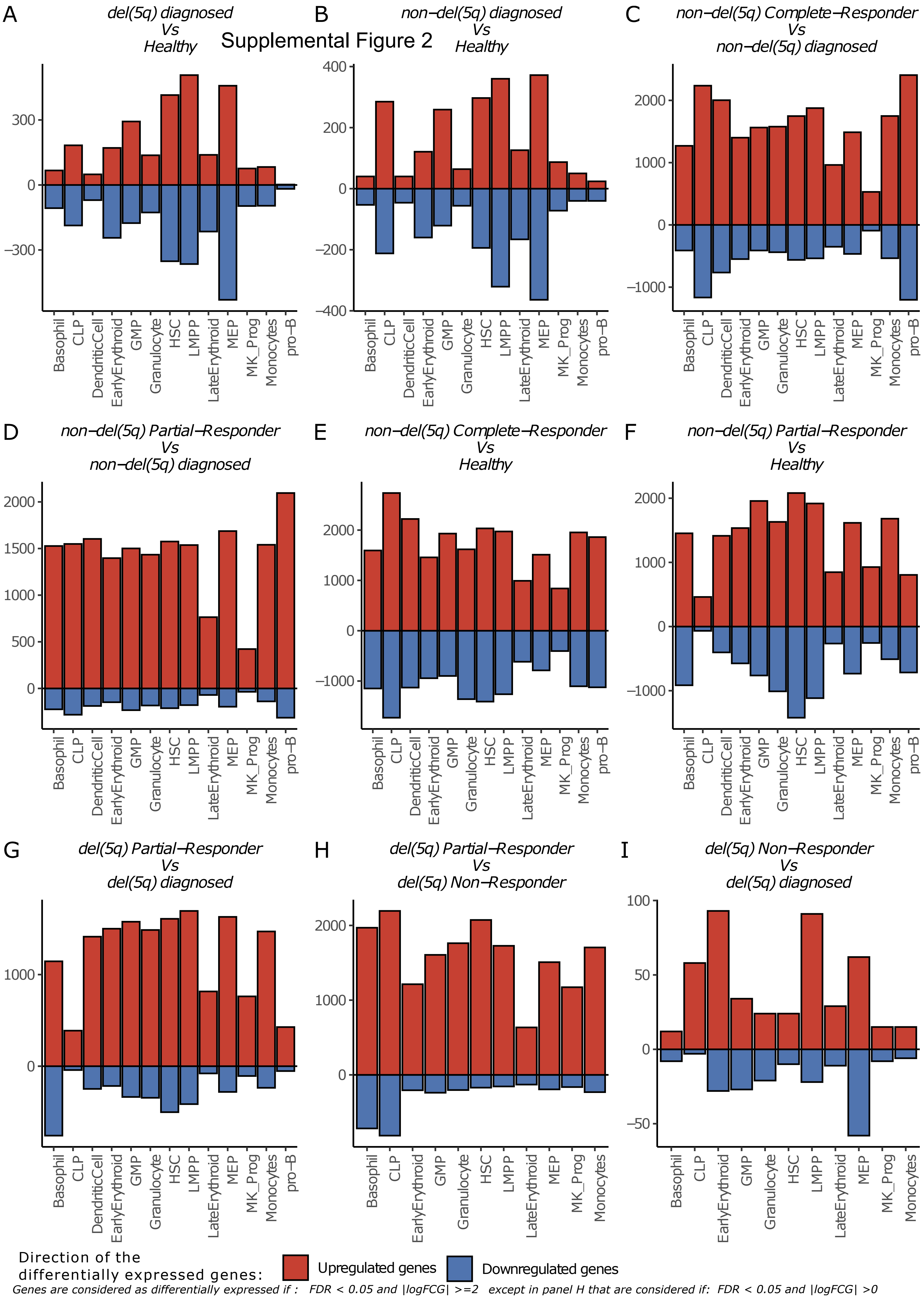


*Genes are considered as differentially expressed if : FDR < 0.05 and |logFCG| >=2 except in panel H that are considered if: FDR < 0.05 and |logFCG| >0*

### Figure S3

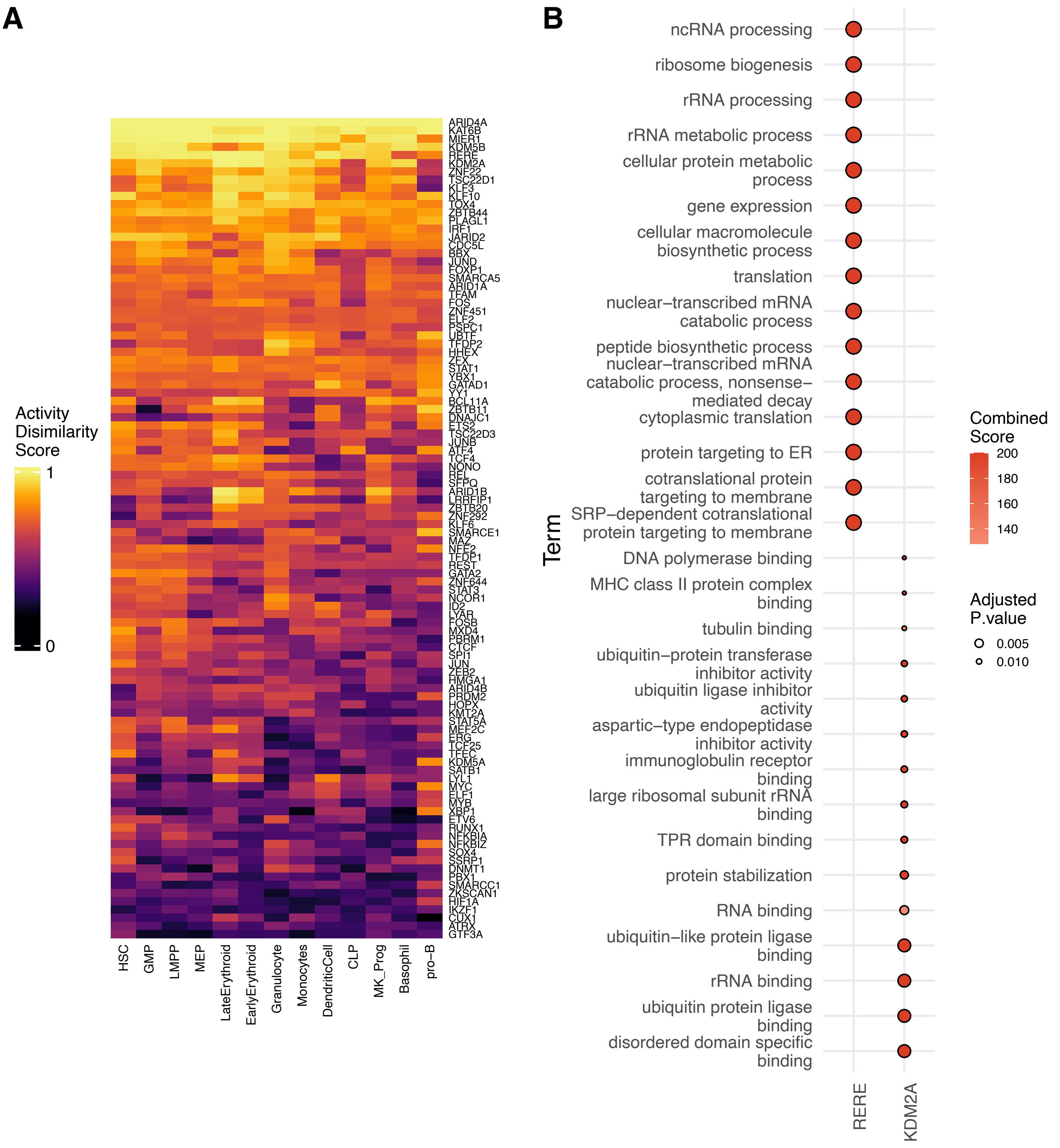

A
B
Activity
Disimilarity
Score
1
0
HSC
GMP
LMPP
MEP
LateErythroid
EarlyErythroid
Granulocyte
Monocytes
DendriticCell
CLP
MK_Prog
Basophil
pro–B
ncRNA processing
ribosome biogenesis
rRNA processing
rRNA metabolic process
cellular protein metabolic process
gene expression
cellular macromolecule biosynthetic process
translation
nuclear–transcribed mRNA catabolic process
peptide biosynthetic process
nuclear–transcribed mRNA catabolic process, nonsense–mediated decay
cytoplasmic translation
protein targeting to ER
cotranslational protein targeting to membrane
SRP–dependent cotranslational protein targeting to membrane
DNA polymerase binding
MHC class II protein complex binding
tubulin binding
ubiquitin–protein transferase inhibitor activity
ubiquitin ligase inhibitor activity
aspartic–type endopeptidase inhibitor activity
immunoglobulin receptor binding
large ribosomal subunit rRNA binding
TPR domain binding
protein stabilization
RNA binding
ubiquitin–like protein ligase binding
rRNA binding
ubiquitin protein ligase binding
disordered domain specific binding
Term
RERE
KDM2A
Combined Score
200
180
160
140
Adjusted P.value
0.005
0.010

### Figure S4

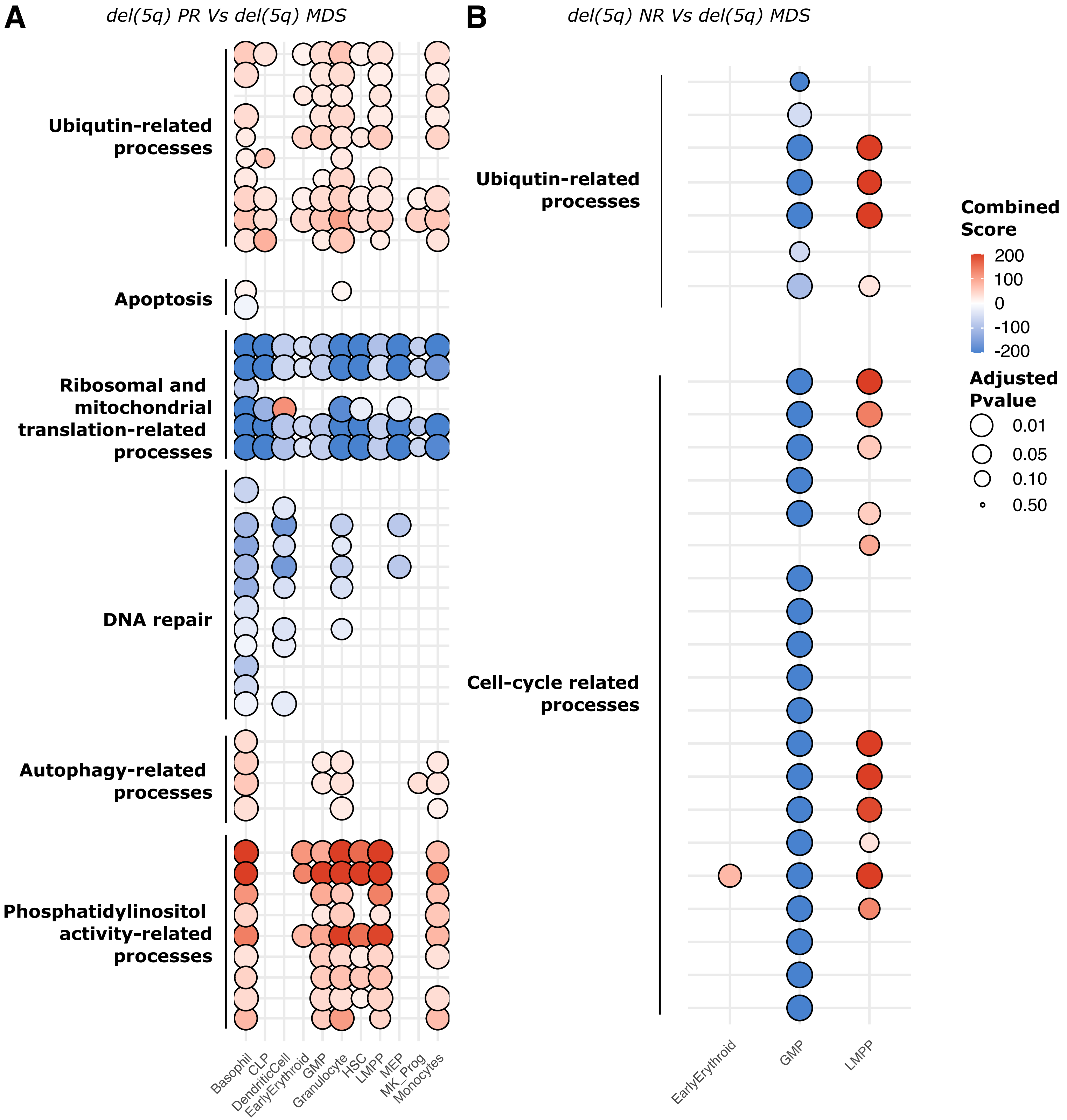

A
del(5q) PR Vs del(5q) MDS
Ubiqutin-related processes
Apoptosis
Ribosomal and mitochondrial translation-related processes
DNA repair
Autophagy-related processes
Phosphatidylinositol activity-related processes
Basophil
CLP
DendriticCell
EarlyErythroid
GMP
Granulocyte
HSC
LMPP
MEP
MK_Prog
Monocytes
B
del(5q) NR Vs del(5q) MDS
Ubiqutin-related processes
Cell-cycle related processes
EarlyErythroid
GMP
LMPP
Combined Score
200
100
0
-100
-200
Adjusted Pvalue
0.01
0.05
0.10
0.50

#### SUPPLEMENTARY FIGURES

**Figure S1**. Hematopoietic $CD34^{+}$ cells from four independent healthy donors were assayed by scRNAseq. **(A)** An overview of the 35897 cells that passed quality control and filtering for the subsequent analysis in this study. Uniform Manifold Approximation and Projection (UMAP) represents the 13 clusters from 3 integrated healthy donors. **(B)** Dotplot showing the percentage and value of the normalized expression of the canonical marker genes used to assign the cell identity to each cluster. **(C)** Independent UMAP for each donor sample colored by cell type. The annotation performed on the integrated assay is transferable to the unique manifold of each sample. **(D)** Barplot representing the contribution of cells from each donor to the different clusters where all the donors, in different proportions, contribute to all clusters except for the pro-B population which has its origin in the healthy donor 3. **(E)** Barplot representing the number of cells assigned to each cell type and the donor of origin for the different cells.

**Figure S2.** The number of the different genes up and downregulated on the different cell types on the different contrasts. **(A)** Del(5q) cells from MDS samples at diagnosis were summarized as pseudobulk and compared to the pseudobulk of the healthy cells from the healthy samples (adjusted pvalue < 0.05 and |logFC>2|). In the same way, the pseudobulks were calculated for the following contrasts: **(B)** non-del(5q) MDS cells at diagnosis compared to the healthy cells (adjusted pvalue < 0.05 and |logFC>2|); non-del(5q) cells from the complete responder **(C)** and the partial responder **(D)** compared to non-del(5q) cells from MDS patients at diagnosis (adjusted pvalue < 0.05 and |logFC>2|); non-del(5q) cells from the complete responder **(E)** and the partial responder **(F)** compared to healthy cells (adjusted pvalue < 0.05 and |logFC>2|); del(5q) cells from the partial responder compared to del(5q) cells at diagnosis (adjusted pvalue < 0.05) **(G)** and compared to del(5q) cells of the non-responder **(H)**; and **(I)** del(5q) cells from the non-responder compared to del(5q) cells at diagnosis (adjusted pvalue < 0.05 and |logFC>2|)**.**

**Figure S3.** Gene regulatory network comparative analysis. **(A)** Heatmap showing the dissimilarity score of the regulons calculated for del(5q) cells, healthy and non-del(5q) cells phenotypes. Warmer color indicates a more different behavior of the regulon between the three states. **(B)** Dotplot showing the over representation analysis of the target genes forming the regulons of KDM2A and RERE.

**Figure S4.** Functional analysis of the differentially expressed genes before and after treatment. Dotplot showing the over representation analysis between the genes differentially expressed before and after the treatment with lenalidomide on the del(5q) cells from the partial cytogenetic responder and the del(5q) cells from a diagnosed MDS contrast **(A)** and the del(5q) cells from the non-responder and the del(5q) cells from a diagnosed MDS contrast **(B).**
